# Soffritto: a deep-learning model for predicting high-resolution replication timing

**DOI:** 10.1101/2025.01.23.634644

**Authors:** Dante Bolzan, Ferhat Ay

## Abstract

**Motivation:** Replication Timing (RT) refers to the order by which DNA loci are replicated during S phase. RT is cell-type specific and implicated in cellular processes including transcription, differentiation, and disease. RT is typically quantified genome-wide using two-fraction assays (e.g., Repli-Seq) which sort cells into early and late S phase fractions followed by DNA sequencing yielding a ratio as the RT signal. While two-fraction RT data is widely available in multiple cell lines, it is limited in its ability to capture high-resolution RT features. To address this, high-resolution Repli-Seq, which quantifies RT across 16 fractions, was developed, but it is costly and technically challenging with very limited data generated to date.

**Results:** Here we developed *Soffritto*, a deep learning model that predicts high-resolution RT data using two-fraction RT data, histone ChIP-seq data, GC content, and gene density as input. Soffritto is composed of a Long Short Term Memory (LSTM) module and a prediction module. The LSTM module learns long- and short-range interactions between genomic bins while the prediction module is composed of a fully connected layer that outputs a 16-fraction probability vector for each bin using the LSTM module’s embeddings as input. By performing both within cell line and cross cell line training and testing for five human and mouse cell lines, we show that Soffritto is able to capture experimental 16-fraction RT signals with high accuracy and the predicted signals allow detection of high-resolution RT patterns.

**Availability:** Soffritto is available at https://github.com/ay-lab/Soffritto.

## Introduction

Replication timing (RT) refers to the process by which cells duplicate DNA during the S phase of the cell cycle. Different regions of the genome replicate at different times in a cell-type specific RT program. While the precise regulation of the RT program remains to be fully elucidated, its pattern has been found to correlate with chromatin structure, histone marks, lamin association and transcriptional activity (Marchal, Sima and Gilbert 2019). The majority of publicly available RT data is produced using 2-fraction, or 2-stage, assays such as Early/Late (E/L) Repli-Seq (Hiratani *et al*. 2008; Marchal *et al*. 2018). In these 2-fraction assays cells are pulse-labeled with BrdU and sorted into early and late S phase fractions. Labeled DNA from both fractions are sequenced and the log2 ratio of early to late coverage is computed for fixed-size bins (e.g., 10kb or 50kb) genome wide. This log2 ratio represents the average time at which each bin (relative to all other bins) is replicated across a cell population. Two-fraction RT assays are limited in their ability to capture replication at high temporal resolution (Marchal *et al*. 2018). To address this limitation, multi-fraction Repli-Seq protocols that sort cells into more than two S phase fractions have been developed, enabling each genomic bin to be characterized by a normalized vector where each value represents the percentage of cells that replicated within the corresponding fraction (Hansen *et al*. 2010). A 16-fraction assay, denoted high-resolution Repli-Seq, is currently the highest resolution multifraction assay. High-resolution Repli-Seq enables the identification of finer scale RT features such as initiation zones (regions of the genome that replicate earlier than surrounding regions and at a constant rate), breakages (small bumps in RT due to replication origin activity within timing transition regions) and biphasically replicating regions that are either undetected or only partially detected in 2-fraction and low-resolution multifraction RT assays (Zhao, Sasaki and Gilbert 2020; Klein *et al*. 2021).

Computational methods have been developed to predict the E/L ratio generated from 2-fraction assays using either epigenomic or sequence-based features. Replicon, a mechanistic model, simulates replication timing using DNase I hypersensitivity, a measure of chromatin accessibility as input (Gindin *et al*. 2014). Another approach based on epigenomic features built LASSO regression models using chromatin binding proteins and histone modifications as input to predict replication timing in *Drosophila* (Comoglio and Paro 2014). TIGER is an algorithm that infers RT profiles from whole genome sequence data obtained from proliferating cell samples by amplifying read depth variation across loci that occur due to differences in RT (Koren, Massey and Bracci 2021). Of the models that utilize DNA sequence, CONCERT, an LSTM-based deep learning model, predicts replication timing by extracting sequence features such as k-mer counts (Yang *et al*. 2022). These methods predict a single value, the average replication timing, per genomic bin, and to date no existing computational approaches predict high-resolution (i.e., 16-fraction) RT. Developing such a predictive model would further our understanding of how exogenous features contribute to finer resolution RT patterns. Furthermore, high-resolution Repli-Seq is a technically challenging and costly assay, thus a predictive model would enable researchers to predict the 16-fraction profiles in cell lines for which such data is not yet available.

With that motivation, we developed Soffritto, a deep learning model that predicts 16-fraction RT from a combination of epigenomic and sequence-based features. More specifically, Soffritto takes six histone modifications (H3K27ac, H3K27me3, H3K36me3, H3K4me1, H3K4me3, H3K9me3), GC content, gene density, and 2-fraction Repli-Seq data as input at 50kb resolution to generate a 16-fraction probability vector for each genomic bin. We validated Soffritto in five cell lines with available 16-fraction RT data, three of which are human-derived (H1, H9, HCT116) and two of which are from mouse (mESC and mNPC). We found that Soffritto accurately captures 16-fraction RT by first training and testing within cell lines and then performing cross-cell line validation. We used five different metrics for evaluation that prioritize different patterns in the 16-fraction RT signal that are useful for its biological interpretation. We also found that Soffritto’s predicted RT profiles captured temporal features specific to high-resolution assays. Overall, this work paves the way for the prediction of high-resolution RT program in distinct cell types to further elucidate the relationship between fine-scale RT patterns and other biological processes.

## Methods

### Data Collection and Processing

The processed 16-fraction Repli-Seq data for H1, H9, HCT116, mESC, and mNPC were downloaded from GSE137764 (Zhao, Sasaki and Gilbert 2020). The data is in the form of a matrix where each column represents a 50kb bin and each row corresponds to an S phase fraction. The rows are sorted from earliest S fraction (S1) to latest S fraction (S16). The data is at 50kb resolution because it is the technical limit for 16-fraction Repli-Seq based on a replication fork speed of 1.8 kb/min and the 30 minute duration of BrdU labeling for each fraction (Zhao, Sasaki and Gilbert 2020). The entries of the matrix are Gaussian-smoothed log2 ratios of the RPKM of the respective S phase fraction to that of a G1 control. Bins with all S phase fraction values of 0 or N/A were removed from downstream analyses. We subsequently normalized each bin such that the 16-fraction values sum to 1. We will refer to these as probability vectors even though each “probability” is a proportion of cells that replicated in the corresponding fraction. These vectors were then used as the labels for model training and testing. We selected six commonly measured histone modifications (H3K27ac, H3K27me3, H3K36me3, H3K4me1, H3K4me3, H3K9me3) for use as features. Histone ChIP-seq data was downloaded from ENCODE (Luo *et al*. 2020) for H1, H9, HCT116, and mESC and from GEO accession GSE96107 for mNPC (Bonev *et al*. 2017). In all cases the histone modification signal was downloaded in bigWig file format and represented as fold-change over control (input). To construct a feature vector for each histone mark, the genome was first partitioned into 50 kb bins. For each bin, the feature’s value was represented as the mean fold change over control.

We downloaded 2-fraction Repli-Seq data for the five cell lines from the 4DN data portal (Reiff *et al*. 2022). We generated 2-fraction RT feature vectors for each cell line by closely following the 4DN Repli-Seq processing pipeline scripts found on https://github.com/4dn-dcic/repli-seq-pipeline. Read alignment files were first merged across biological replicates for early and late fractions respectively. The merged alignments were then filtered, sorted, and deduplicated. Read counts per kilobase million (RPKM) were computed for non-overlapping 50kb bins for early and late fractions respectively. The base 2 logarithm ratio of early to late RPKM was used as the 2-fraction RT feature vector. To avoid division by 0, a small adjustment factor equal to the most frequent difference between consecutive bin RPKM values is added to the early and late RPKMs respectively. All accession IDs for the 2-fraction Repli-Seq data and the histone ChIP-seq data may be found in **Supplementary Table 1**.

Two sequence-based features were utilized: gene density and GC content. To construct gene density feature vectors, we ran *bedtools intersect* to assign a count value to each 50 kb bin that corresponds to the number of overlapping genes with at least one base-pair overlap. Gene coordinates were obtained from GENCODE’s gene annotation (hg38 for human and mm10 for mouse) (Frankish *et al*. 2019). The GC content feature vector was obtained by computing the fraction of base pairs that are either G or C for each 50 kb bin using the hg38 or mm10 reference genome. All nine features were concatenated to form an feature matrix to use as model input where corresponds to the number of genomic bins for which 16-fraction Repli-Seq data was available for a given set of training data.

### Soffritto Architecture

Soffritto consists of a Long Short Term Memory (LSTM) module and a prediction module. LSTMs are a variant of recurrent neural networks that are typically applied to sequential data (Hochreiter and Schmidhuber 1997). Each LSTM cell is composed of an input, forget, and output gate, which coordinate how much information is received from the previous time step (genomic bin in this application). For a given genomic bin *t*, batch size *n*, feature dimension *d*, and hidden dimension *h*, the corresponding LSTM cell is represented by the following equations:

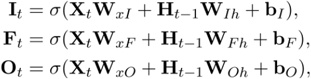

Where **I**_*t*_, **F**_*t*_, **O**_*t*_, ∈ ℝ^*n* ×*h*^ represent the input, forget, and output gates respectively, **X**_*t*_ ∈ ℝ^*n*×*d*^ is the input, **H**_*t*−1_ ∈ ℝ^*n ×h*^ is the hidden state of the previous time step, **w**_*xI*_, **w**_*xF*_, **w**_*xo*_ ∈ ℝ^*d*×*h*^ and **w**_*Ih*_ **w**_*Fh*_ w_*oh*_ ∈ ℝ ^*h*×*h*^ are learnable weight parameters, and **b**_*I*_,**b**_*F*_, **b**_*o*_ ∈ ℝ ^*h*×1^ are biases. The outputs of each gate are then combined to update the cell state and the hidden state of each cell as follows:

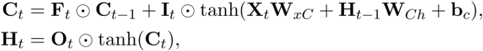

where **C**_*t*_ ∈ ℝ^*n*×*h*^ represents the cell’s internal state, **W**_*x*_*c* ∈ ℝ^*d*×*h*^ and **W**_*ch*_ ∈ ℝ^*h*×*h*^ are weight parameters,b_*c*_ ∈ ℝ^h×1^ is a bias vector, and “circle with the dot” represents the Hadamard product of two matrices (i.e., element-wise multiplication). Soffritto’s LSTM module is bidirectional and therefore an identical set of variables and equations is learned in parallel with recursion being performed in the opposite direction, i.e., the genomic bin to the right rather than to the left. Furthermore, Soffritto’s LSTM module is multilayered with a separate set of parameters learned for each layer while depending on the hidden states of the previous layer. Concretely, given *L* layers and *L* ≥ 2, input 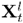 of the *l*-th layer is equal to 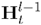, the hidden state of the same bin in the previous layer. In the final layer, each bidirectional 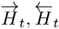 pair is concatenated to form an *n* ×2*h* matrix, which is inputted into Soffritto’s prediction module. The prediction module consists of a fully connected layer that generates a matrix of dimension *n* ×16, where each column corresponds to an S phase fraction, sorted from early to late and each row is a genomic bin. Next, a logsoftmax transformation is applied to each row to facilitate training with KL divergence loss. For prediction, an exponential transformation is applied as a final step to ensure that the RT prediction is in probability space for each bin.

### Model Training

Soffritto was trained using KL divergence loss and L2 regularization to prevent overfitting. The Adam optimization algorithm was used for parameter updates. For intra-cell line training, we left out chromosome 6 for validation and chromosome 9 for testing while the rest of the autosomal chromosomes were used for training. We omitted sex chromosomes to avoid sex bias. For leave-one-cell-line-out (LOCLO) training (cross cell line), we iteratively left out an entire cell line’s data for testing and trained on all autosomes except for chromosomes 6 and 9 from the other four cell lines. The chromosome 6 data from the four cell lines in the training set was used for validation (**Supplementary Figure 1**). All models were trained for 100 epochs. A hyperparameter grid search was performed on the validation chromosome by tweaking the number of LSTM layers (1, 2, 3, 4), the batch size (8, 16, 32, 64), the hidden dimension (16, 32, 64, 128, 256), the learning rate (0.00001, 0.0001, 0.001), and the L2 weight decay (0, 0.00001, 0.0001, 0.001). The model with the lowest KL divergence loss on the validation set was selected to predict on the test set. Soffritto was implemented in PyTorch (Paszke *et al*. 2019).

### Feature-specific Baseline Predictions

For each feature, we first sorted values from lowest to highest and then partitioned into 16 quantile bins. For each bin we assigned an S fraction label based on its order for both ascending and descending values. For example, in ascending order, the quantile bin with the highest values is given a label of S16, but a label of S1 in descending order. We then assigned S fraction labels to each genomic bin according to its quantile label. For evaluating these predictions, we used mean absolute error (MAE) between the predicted S fraction labels and the observed Argmax RT fractions across bins.

### Baseline Predictive Models

Given that there are no existing 16-fraction RT prediction methods, we chose the following three models as baselines to contextualize Soffritto’s performance: support vector regression (SVR), linear regression, and random forest regression. All models were implemented using the *scikit-learn* package. For all three models and for each train-test split, we trained a separate model with default parameters for each S phase fraction. The models’ predicted values were clipped between 0 and 1 and then normalized to sum to one for each genomic bin. The predicted values were clipped prior to normalization to ensure that negative values would not skew the normalization and contradict the probabilistic definition of the labels.

### Soffritto Hidden State Analysis

Soffritto’s hidden states are defined as the embeddings outputted from the last layer of LSTM module for each genomic bin in the test set. Each hidden state vector is of dimension 2*h*, where *h* is the hidden embedding due to concatenation of the forward and reverse LSTM chains. The neighborhood graph was computed using Scanpy’s *neighbors* function with default parameters (number of neighbors = 15) (Wolf, Angerer and Theis 2018). The UMAP (McLnnes, Healy and Melville 2020) embeddings were also computed using Scanpy’s default parameters.

## Results

### Soffritto accurately captures high-resolution RT within cell lines

To evaluate Soffritto, we trained separate models for each of the five cell lines (H1, H9, HCT116, mESC, and mNPC) on all autosomal chromosomes except for chromosome 6 and chromosome 9, which were used for validation and testing respectively. The baseline models were trained and tested on the same data splits. While Soffritto’s predicted 16-fraction RT heatmaps visually recapitulate experimentally measured data to a great extent (**Figure 2A**), we sought to quantitatively assess predictions using five different metrics that capture both the general probabilistic nature of the labels and attributes that are useful for characterizing RT.

**Figure 1:**
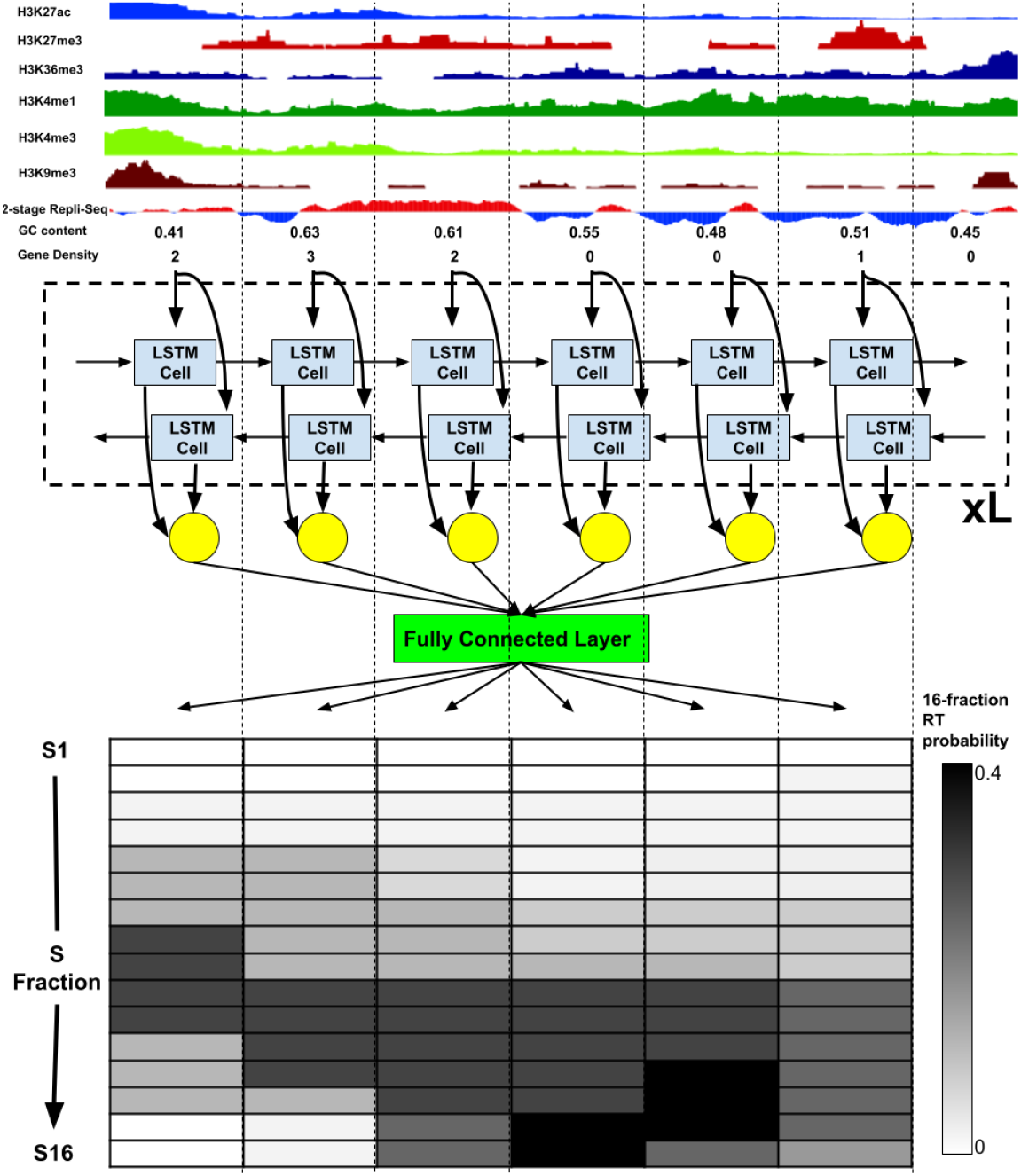
Soffritto architecture. Each vertical dotted line delineates a 50kb bin. Nine features are fed into a bi-directonal, multi-layered LSTM module and a prediction module applies a fully connected layer to generate 16-fraction replication probabilities for each bin.

**Figure 2:**
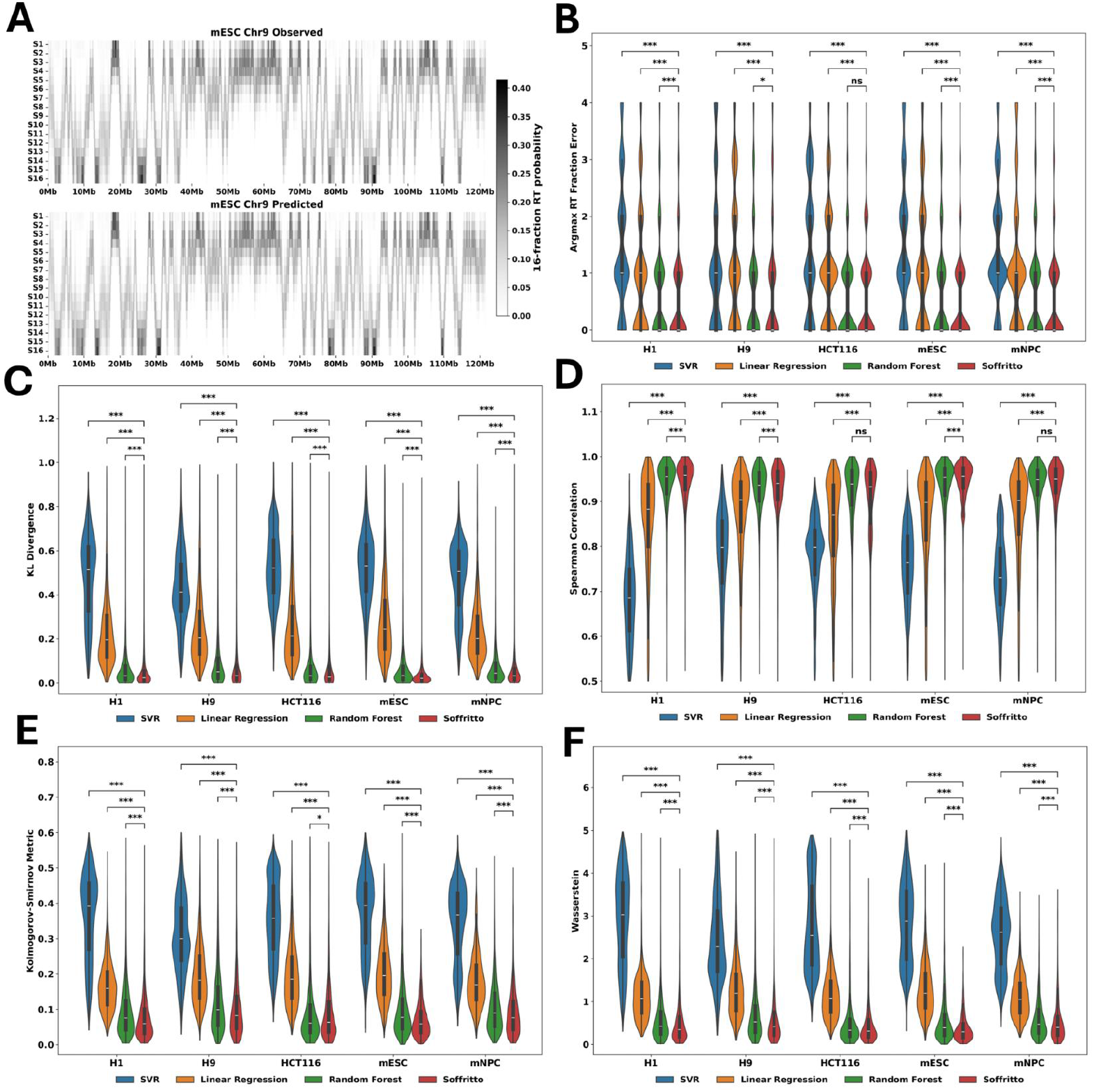
Intra-cell line evaluation of Soffritto on chromosome 9. **A)** Observed vs. predicted 16-fraction Repli-Seq heatmaps in mESC for the whole chromosome 9. **B-F)** Violin plots of the five metrics with each data point corresponding to a bin. All statistical tests were performed using a Wilcoxon signed-rank test between Soffritto’s predictions and each of the baseline models’ predictions. Each Wilcoxon test was one-sided with the alternative hypothesis being that Soffritto’s mean was greater than the baseline model’s mean for Spearman Correlation and less than for all other metrics; ns = not significant (p-value > 0.05), *: 0.01 < p-value < 0.05, **: 0.001 < p-value <= 0.01, ***: p-value <= 0.001

#### Argmax RT Fraction Error

We first examined the S phase fraction which contained the highest replication probability (argmax) for each bin for both observed and predicted RT profiles for chromosome 9. While this argmax RT fraction does not capture cellular heterogeneity in RT, it provides an overall measure of the dominant fraction at which a genomic bin is replicated. We define the metric Argmax RT Fraction Error (ARFE) as the absolute difference between the argmax RT fraction of observed and predicted high-resolution RT data. ARFE yields an integer value that lies between 0 and 15 for each bin (argmax fractions are represented as integers from 0 to 15) with 0 being the best possible agreement. ARFE has the advantage of being biologically interpretable: if a bin has an ARFE value of 2, then the model’s prediction is 2 S phase fractions off from the observed/measured data. We found that Soffritto achieves a mean ARFE of less than one for each cell line (e.g., mean of 0.55 for mESC) as well as a median ARFE of 0 for all cell lines. Compared to the baseline predictive models, Soffritto achieves the lowest mean ARFE significantly in four out of the five cell lines across all bins (**Figure 2B**). We also created baseline models for each input feature by partitioning feature values into 16 quantiles and predicting the corresponding S phase fraction based on ascending or descending order (**Methods, Supplementary Figure 2**). We found that Soffritto’s mean ARFE is consistently lower across all cell lines compared to each feature-specific baseline with 2-stage replication timing being the best predictor among all features (**Supplementary Figure 2**).

#### KL divergence

We next examined how similar the overall predicted 16-fraction probability distribution is to the observed distribution for each bin. To quantify this distance, we computed KL divergence for each bin by setting the observed 16-fraction probabilities as the true probability distribution. KL divergence ranges from 0 to infinity with 0 indicating that the probability distributions are a perfect match. With respect to this metric, Soffritto achieved the lowest mean KL divergence across bins in all cell lines. Notably, Soffritto’s predictions had a lower median KL divergence than the random forest model, with wider and tighter distributions around the median (**Figure 2C**).

#### Spearman correlation

The monotonic order of the S phase fractions in terms of replication probability is important to consider when evaluating a model’s predictions because it is more relevant biologically to capture whether a greater percentage of cells replicate a particular bin for one S phase fraction over another rather than predicting the exact replication probabilities. For this reason and for the interpretable nature of rank correlation, we included Spearman correlation as an additional metric in our analysis. For each bin we compute a Spearman correlation between predicted and observed 16-fraction probability values. Here we found that Soffritto’s predictions highly correlate with observed values with median values ranging from 0.93 to 0.96 across the five cell lines. In addition, Soffritto’s predictions had a significantly higher mean correlation than all baseline models across all cell lines, except in HCT116 and mNPC where there was no significant difference between predictions of Soffritto and Random Forest model (**Figure 2D**).

#### Kolmogorov-Smirnov and Wasserstein Distance

and Another way of conceptualizing 16-fraction RT is by examining the cumulative replication fraction (Zhao, Sasaki and Gilbert 2020). The cumulative replication fraction of a specific S phase fraction is defined as the cumulative sum of replication probabilities for all earlier fractions and the current fraction. Consequently, we obtain a vector length 16 for each bin that now represents a cumulative distribution rather than a probability distribution. In this space, one can infer the percentage of cells that replicated a genomic bin by a specific S phase fraction. We computed two metrics that can both be formulated as a measure of distance between two cumulative distribution functions (CDF): Kolmogorov-Smirnov (Massey 1951) and Wasserstein Distance (Kantorovich 1960). The Kolmogorov-Smirnov statistic is defined as the maximum absolute difference between empirical CDFs of two samples while the Wasserstein Distance implicitly computes the sum of the absolute differences between two CDFs. We chose both metrics because they capture both overall similarity between two CDFs and penalize the greatest error between two CDFs. We observed that Soffritto achieves a significantly lower mean Kolmogorov-Smirnov value than the baseline models in all cell lines, including in HCT116, for which Soffritto and Random Forest have performed similarly with respect to non-CDF metrics (**Figure 2E**). As for Wasserstein Distance which captures a more global distance between CDFs, Soffritto achieves both a lower mean and median than all baselines in all cell lines, with particularly tighter distributions for the mouse cell lines (**Figure 2F**). To quantify Soffritto’s overall performance, we ranked the models based on mean values for each of the five metrics and the five cell lines for a total of 25 rankings. We then computed the Borda count (Emerson 2013) for each ranking (0 points for the lowest-ranked model, 1 point for the second-lowest ranked model, etc.) and summed the points for each model across all rankings. Soffritto was the highest ranked model with 74 out of a possible 75 points. Overall these results demonstrate that Soffritto’s predictions capture important RT characteristics in both probability space and CDF space.

### Soffritto’s predicted RT profiles capture RT dynamics

Next we aimed to examine whether Soffritto’s intra-cell line predictions captured high-resolution RT dynamics in the test chromosome. Two important features of RT dynamics that capture RT heterogeneity are *Trep* and *Twidth* (Zhao, Sasaki and Gilbert 2020). *Trep* is defined as the time point at which 50% of cells have finished replicating a genomic bin while *Twidth* is defined as the difference in time between when 75% of cells have finished replicating and 25% of cells have finished replicating. To compute *Trep* and *Twidth*, a sigmoidal curve is fit to the cumulative replication fraction values for a particular bin (**Figure 3A**). For our analysis, we express time in continuous values ranging from 0 (S1) to 15 (S16). We computed *Trep* for both predicted and observed 16-fraction data for each bin in chromosome 9 for all five cell lines, and we found that Soffritto’s predicted *Trep* values correlate highly with observed *Trep* values, with a mean correlation of 0.96 (median 0.97) (**Figure 3B**).

**Figure 3:**
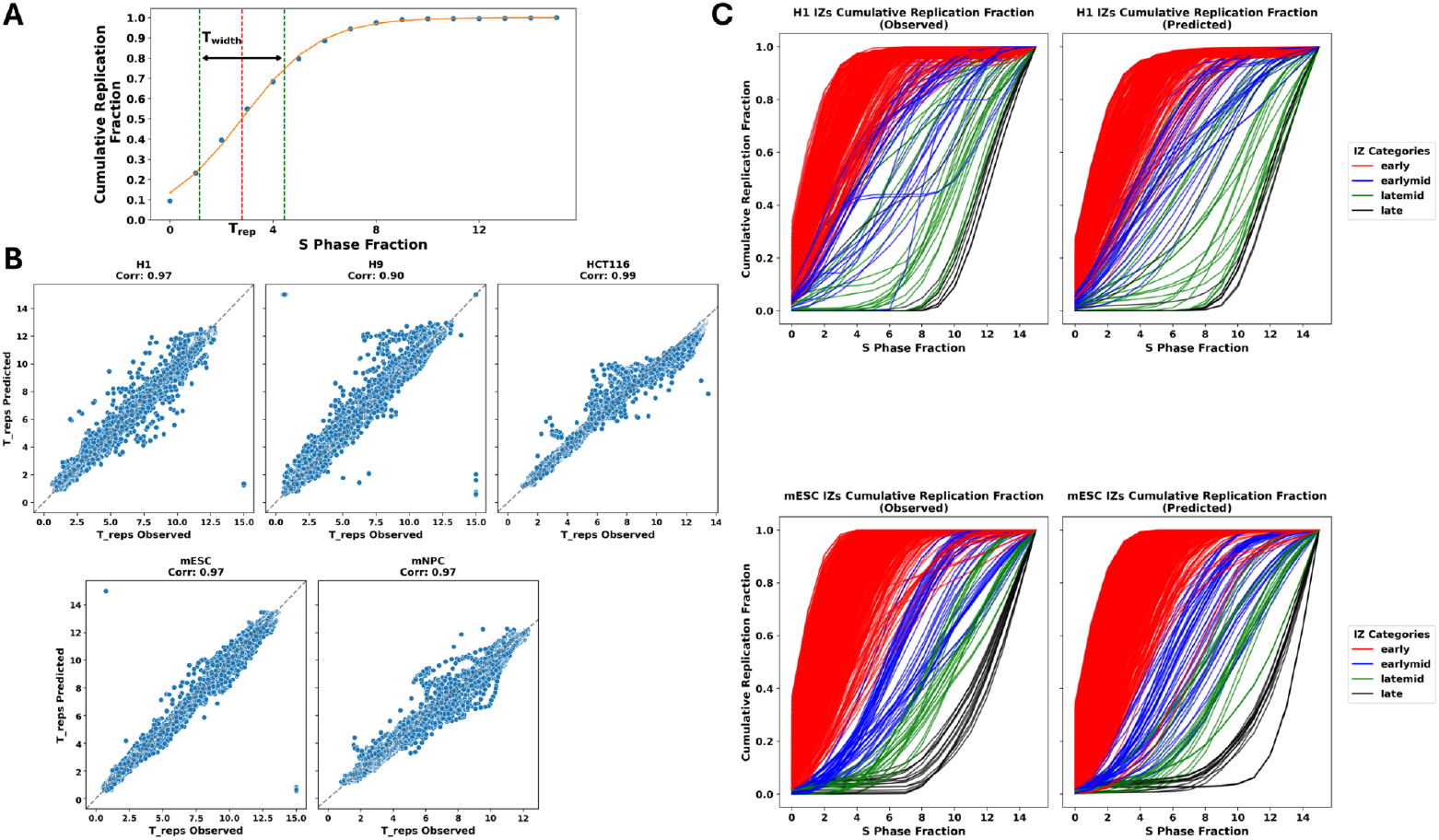
Dynamic RT features for observed vs. predicted RT profiles. **A)** Demonstration of *T*_*rep*_ and *T*_*width*_ calculation using a representative cumulative replication fraction distribution of a single bin. **B)** Pearson correlation plots for observed vs. predicted *T*_*rep*_ in all cell lines with each data point corresponding to a single bin of chromosome 9. The y-axis and x-axis correspond to predicted and observed *T*_*rep*_ values respectively. **C)** Observed vs. predicted cumulative replication fraction curves grouped by temporal category in bins classified as Initiation Zones (IZs) for H1 (top) and mESC (bottom). Each curve corresponds to a single bin of chromosome 9 that is categorized as IZ by Zhao et al., 2020.

We then compared the replication dynamics of initiation zones for both observed and predicted 16-fraction RT to determine if Soffritto is capturing expected dynamics at these sites. Initiation zones (IZs) are defined as consecutive genomic bins that replicate earlier than flanking regions and at a roughly constant rate. They are thought to initiate replication as they are distributed throughout the genome and RT does not occur in a uniform manner genome-wide. In addition, IZs can be classified into four temporal categories based on their S phase fractions in which highest read density was identified: early (S1-S3), early-mid (S4-S6), late-mid (S7-S9), and late (S10-S12) (Zhao, Sasaki and Gilbert 2020). We obtained the genomic coordinates for IZs along with their corresponding temporal labels in H1 and mESC from (Zhao, Sasaki and Gilbert 2020). We then computed the cumulative replication fraction for all bins located in IZs for both observed and predicted 16-fraction RT profiles on chromosome 9. We found that Soffritto’s predictions capture the increase in *Twidth* as S phase progresses from early to mid, followed by a decrease in *Twidth* from mid-S phase to late S phase in both H1 and mESC (**Figure 3C**). These findings suggest that Soffritto’s predictions capture high-resolution temporal features that can be measured specifically in 16-fraction RT data.

### Soffritto’s hidden states reveal early to late RT transitions and highlights outlier regions

In order to better interpret and understand Soffritto’s predictions, we next extracted Soffritto’s hidden states for downstream analysis. More concretely, for each cell line, we passed the test set (chromosome 9) through the corresponding trained Soffritto model’s LSTM module to obtain the hidden state matrix of the last layer of the LSTM module. The hidden state matrix is an *n* x 2*h* matrix, where *n* is the number of bins in the test chromosome and *h* is the hidden dimension (**Methods**). We then computed a nearest neighbor graph using this hidden state matrix as input and then embedded it using UMAP. **Figure 4** shows the UMAP plots for all five cell lines with each dot corresponding to a 50kb bin on chromosome 9. We annotate each genomic bin according to its Argmax RT Fraction to characterize that bin’s general RT (**Figure 4A-E**). We observe that Soffritto’s hidden states capture broad early to late RT transitions across bins for all cell lines, suggesting that Soffritto learns general RT before performing precise 16-fraction predictions.

**Figure 4:**
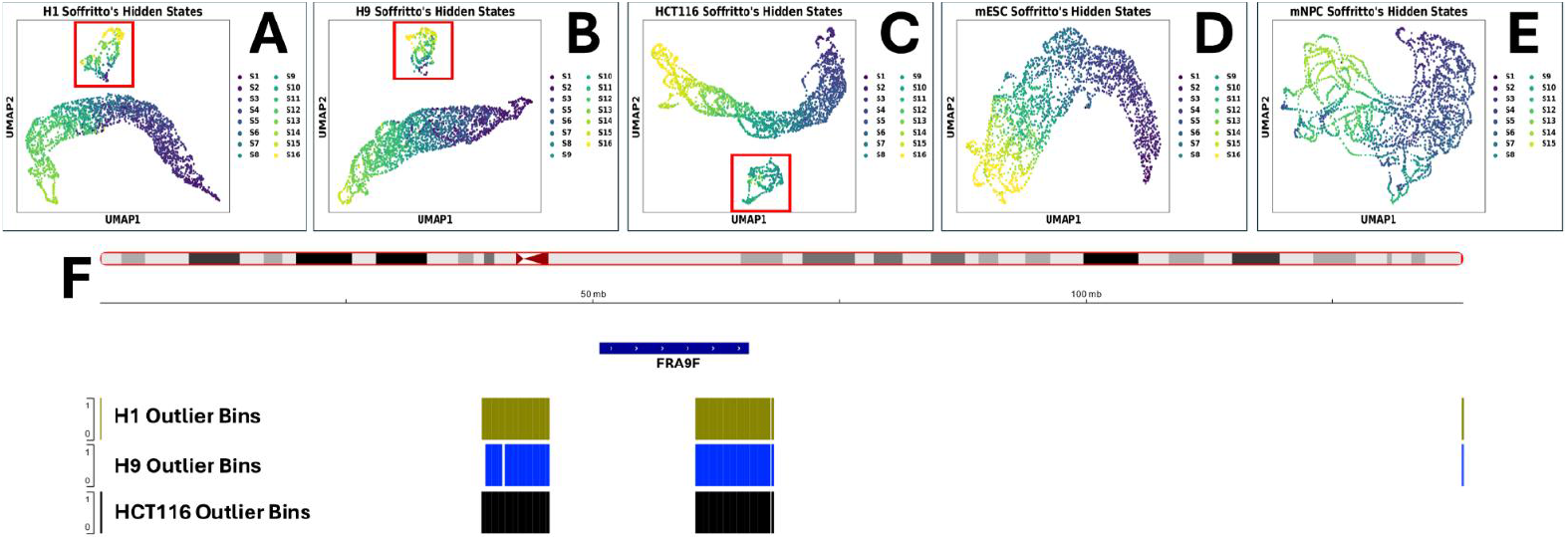
Soffritto hidden state plots. **A-E)** UMAP plots generated from Soffritto’s hidden state embeddings for chromosome 9 in H1, H9, HCT116, mESC, and mNPC. Each dot corresponds to a genomic bin. Each bin is annotated with its Argmax RT Fraction (S1-S16), **F)** Genomic tracks for the entire chromosome 9 (hg38 build). The track at the top shows the coordinates of FRA9F, a common fragile site. The next three tracks display the coordinates for the genomic bins enclosed by the red rectangles in UMAP plots A-C for H1, H9, and HCT116 respectively.

In addition, for the three human cell lines, a group of bins were separated from the rest in UMAP space, which we highlight with red rectangles (**Figure 4A-C**). While bins in these clusters tend towards late replication, they show a broad spectrum of RT values similar to the bins not present in these clusters, suggesting that their separation is not driven by their RT profiles. All of the clusters have roughly the same cardinality across the three cell lines: 296, 277, and 295 bins for H1, H9, and HCT116 respectively. We then obtained the genomic coordinates of these bins and found that 274 bins are shared among the three cell lines. The high number of intersecting bins suggest that sequence-based features are driving their separation in the UMAP plot. We found that the vast majority of bins localize into two consecutive blocks (**Figure 4F**), one spanning the entire chromosome 9 centromere and the other overlapping with FRA9F, the largest common fragile site on chromosome 9 (Kumar *et al*. 2019). Centromeric DNA is highly repetitive and compact, making it susceptible to DNA replication stress, defined as the slowing or stalling of DNA replication (Carnie *et al*. 2023). Furthermore, 16-fraction RT data is generated using short reads making it difficult to measure precise RT in repetitive regions such as centromeres. Fragile sites are defined as loci prone to gaps and breakages when DNA replication is partially inhibited (Schwartz, Zlotorynski and Kerem 2006; Li and Wu 2020). The remaining bins were localized near the p-arm telomere in H1 and HCT116 and adjacent to the q-arm telomere in H1 and H9 (**Figure 4F**). We also obtained the hidden states for chromosomes in the training set and also observed a small set of bins that separated from the rest in UMAP space in the human cell lines for chromosomes 2, 18, and 22 (**Supplementary Figure 3**). These clusters tended to be more late replicating than the rest of the genome, but exhibited RT heterogeneity both within and across cell lines. While these clusters did not overlap with any fragile sites, they consistently formed continuous chunks overlapping the centromere or adjacent regions (**Supplementary Figure 4**). For all chromosomes mentioned, these clusters were also found to have significantly lower histone modification signals (for all six histones that were utilized as features) than the other genomic bins (**Supplementary Figure 5**). Given that these outlier regions lack both active (e.g., H3K27ac) and repressive marks (e.g., H3K9me3), they are reminiscent of Quiescent states defined by ChromHMM (Ernst and Kellis 2012). Overall, these results suggest that the hidden states learned by Soffritto capture a continuous spectrum for ordering RT fractions and identify genomics regions (outliers) for which the relationship between RT and histone marks deviate from this spectrum.

### Soffritto’s predictions generalize to unseen cell lines

We next wanted to test whether Soffritto could predict 16-fraction RT in an unseen cell line when trained on data from multiple other cell lines. To do this we implemented a leave-one-cell-line-out strategy by iteratively leaving one cell line out while training on the other four cell lines (**Methods**). We omitted chromosome 9 from the training data to ensure an unseen chromosome in the test set to account for potential model memorization from the sequence-based features (with the caveat that there is no synteny between human chr9 and mouse chr9). We observed that Soffritto’s predicted RT captured the general RT trends of chromosome 9 to a good extent (**Figure 5A;** median Spearman correlation of 0.91). For a subset of regions (e.g., the last 20 Mb of chromosome 9), the predicted RT showed a wider distribution across adjacent fractions compared to observed profiles (**Figure 5A**).

**Figure 5:**
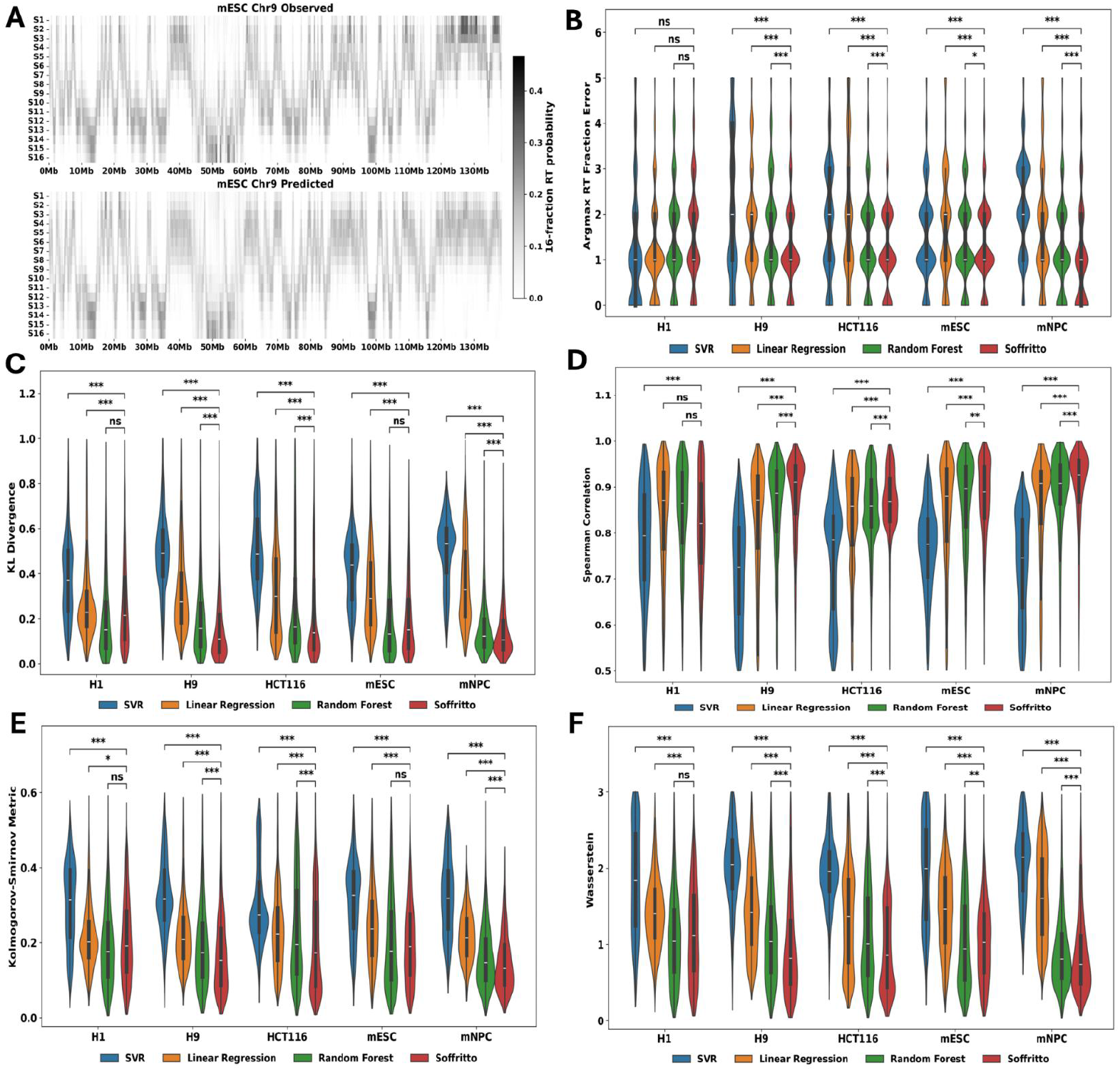
Leave-one-cell-line-out (cross cell line) results. **A)** Observed vs. predicted RT heatmaps for left out H9. **B-F)** Each panel corresponds to a different metric. Each cell line on the x-axis corresponds to the cell line left out when training the model. All p-values were computed and annotated with symbols as was described in Figure 2.

We also trained the same three general baseline models (SVR, linear regression, random forest) as in the intra-cell line evaluation on this cross-cell line setting. We evaluated Soffritto using the same five metrics as the intra-cell line analysis. Soffritto achieved a significantly lower mean ARFE than the baseline models for four of the five left out cell lines, being on average 1.4 S phase fractions off (**Figure 5B**). Figure 5C shows that Soffritto has a significantly lower mean KL divergence for H9, HCT116, and mNPC with comparable performance between Soffritto and random forest in mESC (**Figure 5C**). Soffritto also achieved a mean Spearman correlation of 0.83 across all left out cell lines, higher than the baseline models for all cell lines except for H1 (**Figure 5D**). With respect to the cumulative replication fraction metrics, Soffritto’s predictions do slightly better according to Wasserstein distance with predictions from Soffritto and random forest for mESC having similar distributions (**Figure 5E-F**). In terms of Soffritto’s overall performance, we ranked the models by computing the Borda count for each ranking and summing the points for each model across all rankings. Within cell lines there are 15 possible points and we found that Soffritto was the highest ranked model for four out of five left out cell lines. Across all cell lines combined, Soffritto gathered 67 out of 75 possible points and random forest, the second highest performing model, got 53 points. Overall, these results demonstrate that Soffritto is able to predict 16-fraction RT patterns in unseen cell lines.

## Discussion

We introduced Soffritto, an LSTM-based model that predicts 16-fraction RT data using two-fraction RT, six histone modifications, GC content, and gene density as input. As the first predictive model of 16-fraction RT, Soffritto outperforms non-recurrent baseline models in intra-cell line and cross-cell line experiments in five human and mouse cell lines. The five metrics we used for evaluation capture different aspects of the 16-fraction RT signal, such as the fraction with the highest probability and the cumulative replication fraction distribution, that are biologically meaningful. We also showed that Soffritto accurately captures patterns of dynamic RT features defined in experimental 16-fraction RT data. Soffritto’s embeddings capture broad RT trends across bins as well as centromeres, which are known for having unstable replication profiles.

Although Soffrito’s predictions for unseen cell types had reasonable accuracy, an increase in the availability of high-resolution (16 or similar number of fractions) RT data could provide substantial benefit to its generalizability. In this work, we used a small set of features to make Soffritto applicable to a broader set of cell types; however, our model can certainly be expanded to include more features either from DNA sequence or cell-type-specific functional genomics measurements. We also note that while Soffritto was the best performing model overall, random forest had comparable performance for multiple train-test splits and metrics suggesting that the input features for each bin have high predictive power even when considered in isolation from neighboring bins. Future improvements for Soffrito may include utilization of input features at higher resolution and/or exploring more sophisticated deep learning architectures that may exploit longer-range correlations of input features or physical proximity of linearly distant bins in 3D space.

