## Supplementary Figure for "Soffritto: a deep-learning model for predicting high-resolution replication timing"

**Supplementary Table 1**: Data sources of non-sequence based features. Histone ChIP-seq data was downloaded from ENCODE with the exception of mNPC. The 2-stage Repli-Seq data was downloaded from the 4DN Portal.

| | **H1** | **H9** | **HCT116** | **mESC** | **mNPC** |
|---|---|---|---|---|---|
| **H3K27ac** | ENCFF423TVA | ENCFF988WEQ | ENCFF169MCH | ENCFF163SBS | GSE96107 |
| **H3K27me3** | ENCFF380KPI | ENCFF077DRH | ENCFF717ZKL | ENCFF182FTP | GSE96107 |
| **H3K36me3** | ENCFF370UNF | ENCFF565AYJ | ENCFF024LGD | ENCFF097KTK | GSE96107 |
| **H3K4me1** | ENCFF396RXV | ENCFF142KLG | ENCFF337BPL | ENCFF280RYX | GSE96107 |
| **H3K4me3** | ENCFF730VVX | ENCFF462TXF | ENCFF649ZLF | ENCFF719IZR | GSE96107 |
| **H3K9me3** | ENCFF350QBK | ENCFF891UKM | ENCFF989AAM | ENCFF519ZAT | GSE96107 |
| **2-stage Repli-Seq** | 4DNESEJCRVNR | 4DNESDB2JQ5S | 4DNESXPNEE4Q | 4DNES3BNI8G3 | 4DNESSLDL428 |

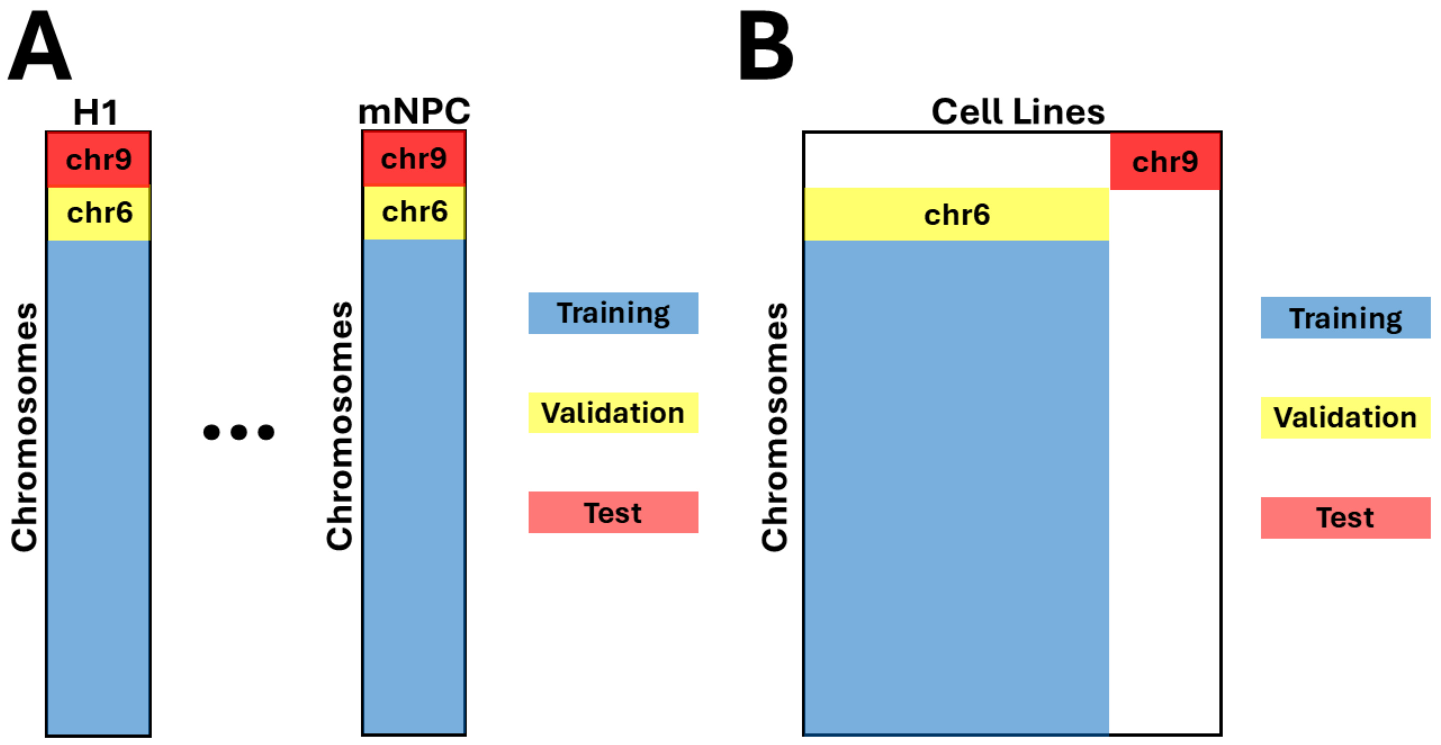


**Supplementary Figure 1**: Train-test splits for intra-cell line (**A**) and leave-one-cell-line-out (**B**) schemes.

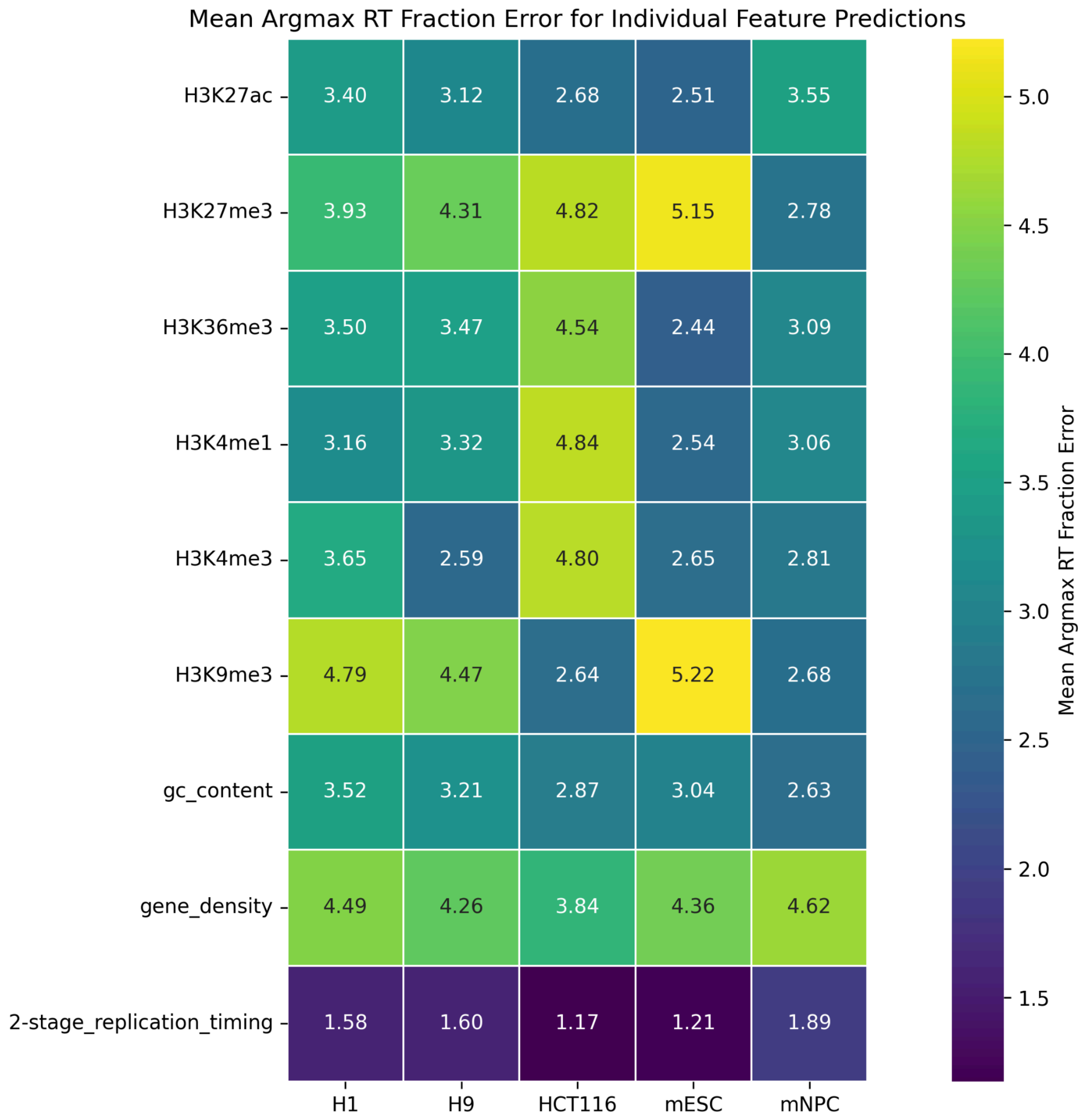


**Supplementary Figure 2**: Mean Argmax RT Fraction Error when predicting from individual features for chromosome 9 for each cell line.The values for each feature were sorted from lowest to highest and then partitioned into 16 non-overlapping intervals. Each interval was assigned a fraction S1-S16 based on ascending or descending order. Each genomic bin was then assigned a fraction based on the interval its feature value fell into. The mean absolute error (chromosome-wise) of these predictions was then computed with respect to the Argmax RT fractions of the observed data for both ascending and descending sorting. The min Argmax RT Fraction Error over both sortings is reported here.

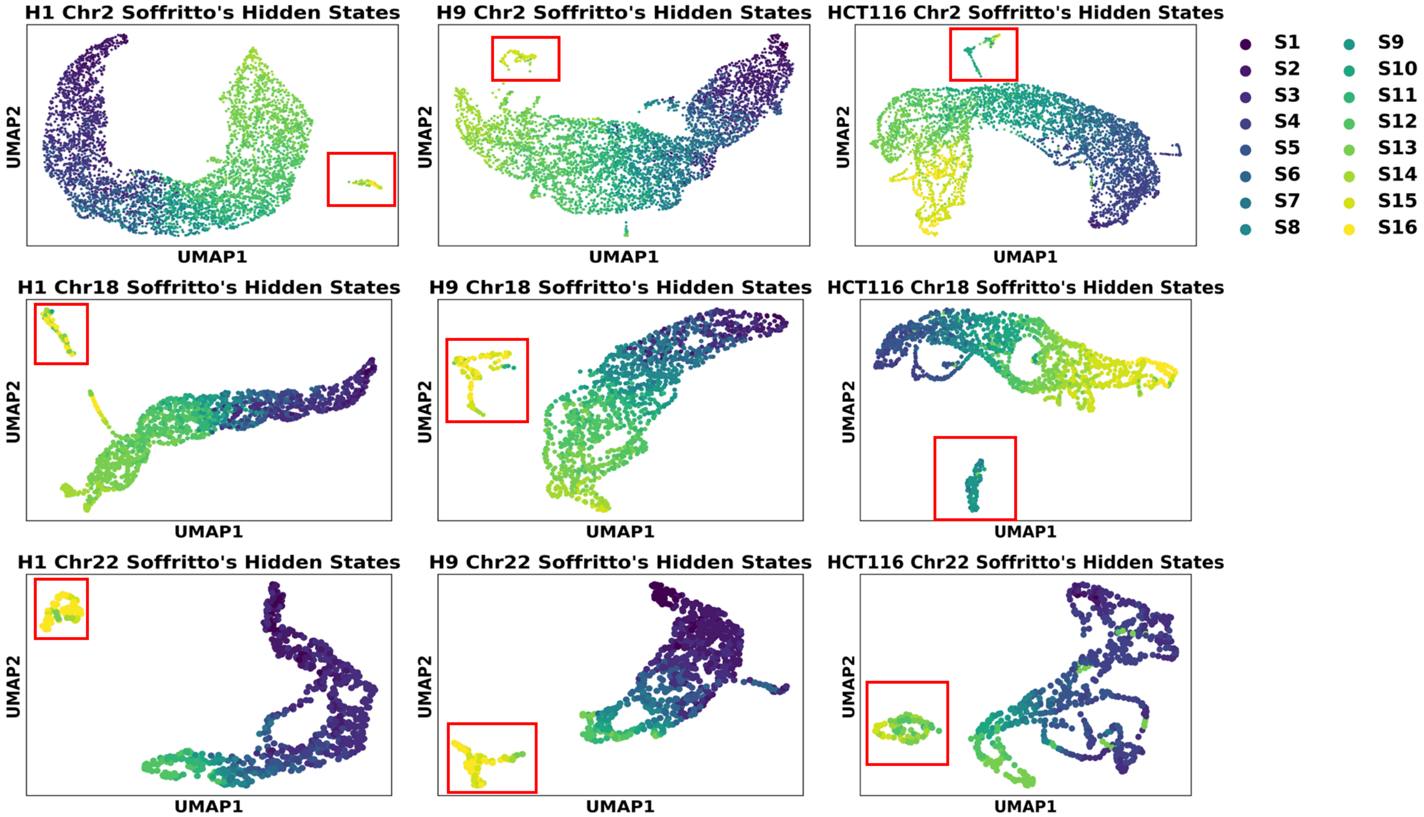


**Supplementary Figure 3**: UMAP plots of Soffritto's hidden states for chromosomes 2, 18, and 22. Each row corresponds to a chromosome and each column a cell line. Outlier bins are outlined with a red rectangle. UMAPs are colored by observed Argmax RT Fraction labels (S1-S16).

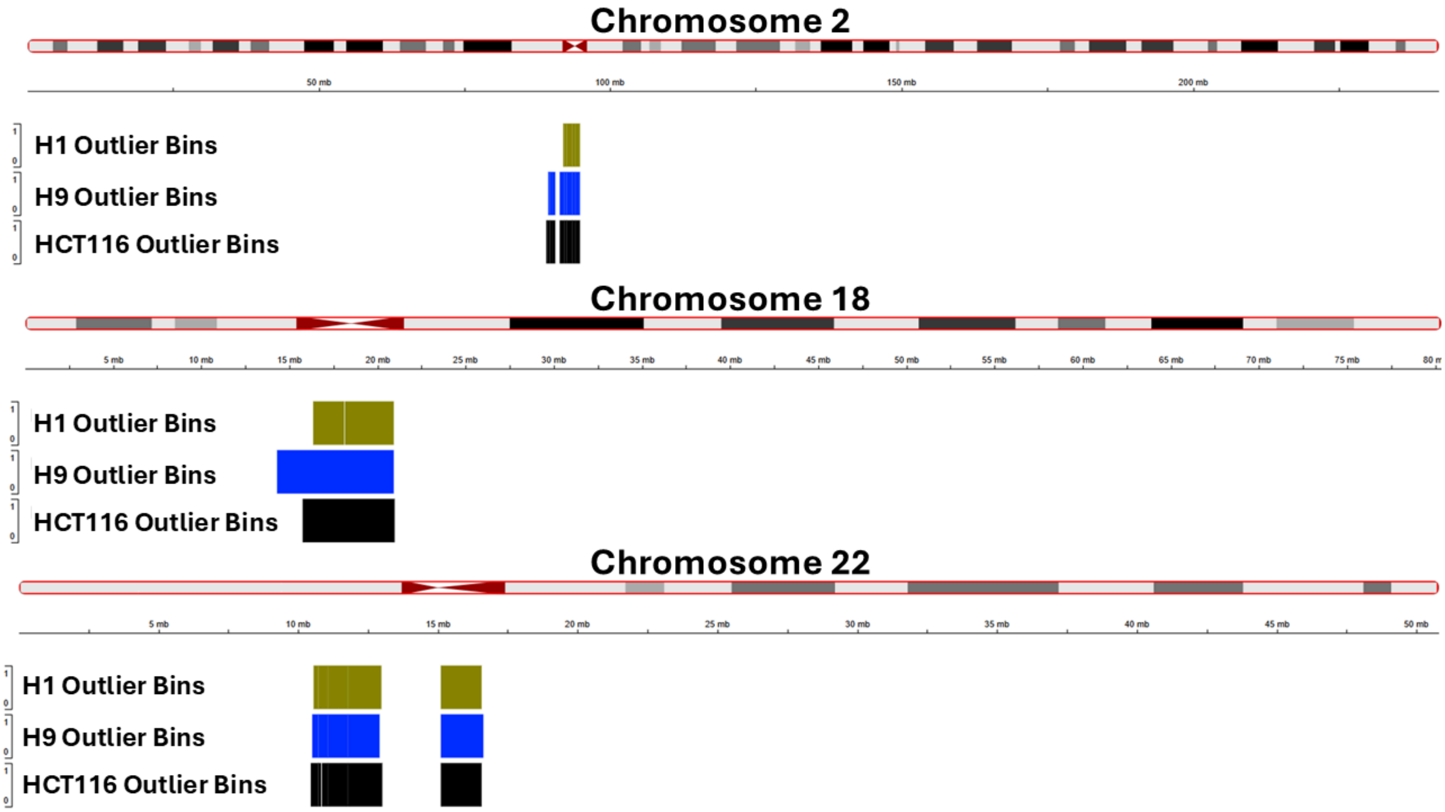


**Supplementary Figure 4**: Genomic tracks of outlying bins identified in Supplementary Figure 3 for chromosomes 2 (top), 18 (middle), and 22 (bottom).

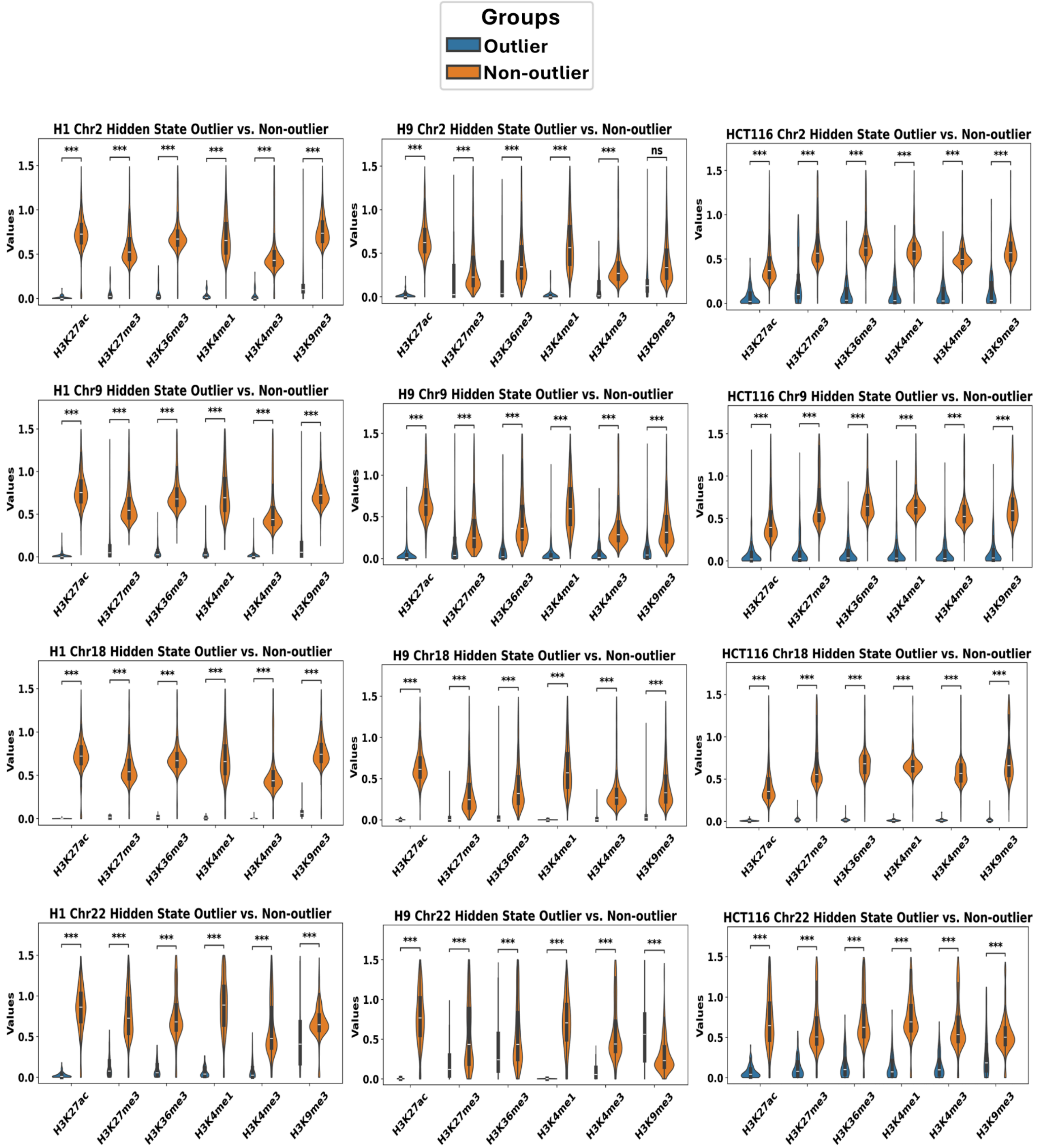


**Supplementary Figure 5**: Violin plots of distributions of histone signal for outlying and non-outlying bins identified by UMAPs of Soffritto's hidden states for chromosomes 2, 9, 18, and 22. P-values are derived from Mann-Whitney U tests; ns = not significant (p-value > 0.05), *: 0.01 < p-value < 0.05, **: 0.001 < p-value <= 0.01, ***: p-value <= 0.001
